# Comparison of a GC-Orbitrap-MS with Parallel GC-FID Capabilities for Metabolomics of Human Serum

**DOI:** 10.1101/740795

**Authors:** Biswapriya B. Misra, Ekong Bassey, Michael Olivier

## Abstract

Gas chromatography mass spectrometry (GC-MS) platforms for use in high throughput and discovery metabolomics have heavily relied on time of flight (ToF), and low resolution quadrupole and ion trap mass spectrometers and are typically run in electron ionization (EI) modes for matching spectral libraries. Traditionally, detectors such as flame ionization detection (FID), have also helped in identification and quantification of compounds in complex samples for diverse clinical applications, i.e., fatty acids. We probed if combination of FID in line with a high-resolution instrument like a GC-Orbitrap-MS may confer advantages over traditional mass spectrometry using EI.

We used a commercially available human serum sample to enhance the chemical space of serum using an advanced high resolution mass spectrometry (HR-MS) platform (QExactive Orbitrap-MS) with an FID feature for confident metabolite identification to assess the suitability of the platform for routine clinical metabolomics research. Using the EI mode, we quantified 294 metabolites in human serum using GC-Orbitrap-MS. These metabolites belonged to 89 biological pathways in KEGG. Following a sample split, using an in-line FID analysis, 1117 peaks were quantified. Moreover, representative peaks from FID and their corresponding MS counterparts showed a good correspondence when compared for relative abundance.

Our study highlights the benefits of the use of a higher mass accuracy instrument for untargeted GC-MS-based metabolomics not only with EI mode but also orthogonal detection method such as FID, for robust and orthogonal quantification, in future studies addressing complex biological samples in clinical set ups.

## 1. Introduction

Metabolomics is the comprehensive study and systematic quantification of small molecules in the molecular weight range of 50-2000 Daltons in biological samples (cells, tissues, organs, biofluids, or whole organisms), and thus, complements efforts from other high throughput omics platforms such as genomics, transcriptomics, and proteomics as an indispensable platform. Platforms for generating metabolomics data typically include gas and liquid chromatography, or capillary electrophoresis linked with mass-spectrometry (GC-MS, LC-MS, and CE-MS), and spectroscopy approaches [such as nuclear magnetic resonance (NMR), infrared (IR), Raman] that have helped address diverse biological questions allowing to connect the genotype with molecular phenotype (1). Particularly, gas chromatography mass spectrometry (GC-MS) is very amenable to polar primary metabolites (such as sugars, amino acids, amines, sugar phosphates, or sugar alcohols) (2) and fatty acids, in addition to excellent chromatographic resolution, thus lending itself to routine quantitative metabolomic applications (2). Newer high resolution (HR) instruments such as Orbitrap mass spectrometers are capable of providing sub-ppm mass accuracy at high mass resolutions (i.e., > 60,000), and hence allow calculation of predicted molecular formulas based on the mass defect of a detected metabolite ion (3–5), and generate mass spectral data at high resolving power with mass accuracies <1 ppm· However, these HRGC-MS platforms have found limited applications till date, baring handful applications in microbial metabolomics (6) and a recent use in non-human primate biofluid (i.e., baboon) serum metabolomics (7).

Most studies have used (GC)-high resolution accurate mass (HRAM) mass spectrometers only in electron ionization (EI) mode of operation. However, flame ionization detection (FID) is without a doubt the most often used gas chromatography (GC) detection method, a technology which dates back to early 1960s and finds applications ranging from analysis of hydrocarbons to fatty acids. When a full spectrum is recorded using mass selective detector (MSD) during a chromatographic run, sensitivity is often inferior in a MS detector when compared to a FID (8). A combination of GC-MS and gas chromatography-flame-ionization detection (GC-FID) also is an old idea, typically run independently and/ or in parallel, with its roots going back to the 1960s. Then the chromatograms are manually aligned and then peaks were partitioned into bins according to retention time values. Unfortunately, comparisons between chromatograms by MS and by GC detectors are difficult since the results vary depending on the samples and the carbon atoms in the molecules being analyzed. FID is sensitive for compounds containing carbon, and its sensitivity is better than thermal conductivity detector (TCD). Previous studies have claimed that GC-FID is considered to be more reliable and sensitive for quantitative analysis than GC–MS, while GC–MS can provide more definite qualitative information and biomolecule identification (9). GC-FID is also considered more sensitive, more reproducible and covers a wider dynamic range when compared to GC-MS in full scan monitoring mode (10).

Recent studies have applied GC-FID in cataloging the human serum metabolomes as a complimentary technique to GC-MS, LC-MS and NMR (11). However, GC-FID and GC-MS or LC-MS as parallel methods have only been used in analysis of bacterial metabolites (12), for targeted fatty acid and lipid characterization in human plasma (13), fecal volatile characterization (14), and transgenic rice metabolism (15) among others. However, none of these analyses were performed using HRMS equipped with both FID and MS detectors that used the same samples at the same time. Combined TLC/GC-FID analysis when compared to GC-MS as the two methods for analysis of human serum lipids, allowed identification and quantification of only eight metabolites in common (arachidonic acid, eicosanoic acid, linoleic acid, oleic acid, palmitelaidic acid, palmitic acid, stearic acid and tetradecanoic acid) (11), suggesting significant complementarity of FID and MS analysis of the human metabolome. Previously, ethanol, methanol, and formate concentrations were measured by headspace GC-FID analysis in vitreous and blood samples collected postmortem (16).

As can be seen, most of these efforts used GC-FID and GC-MS as two independent approaches one after another and not in-line. To our knowledge, studies have not attempted to characterize the complex biological matrixes of clinically relevant samples such as human serum, and to show their joint application in clinical metabolomics, and rather have only been used for targeted chemical constituents such as drugs, pesticides, and organic exogenous chemicals. Our study is the first attempt to identify and quantify serum metabolites using a high mass resolution gas chromatography mass spectrometer (GC-Orbitrap-MS) with two detectors (FID and MS) on a comparative basis. The principal aims of this study were to assess the capabilities of GC-FID analysis in parallel to a GC-Orbitrap-MS analysis for quantification and identification of metabolites in human serum as a test sample, in order to exploit the full capabilities of these two detectors and instrument for untargeted clinical metabolomics.

## 2. Materials and Methods

### 2.1 Chemicals

Solvents such as acetonitrile, isopropanol, and pyridine were of HPLC grade, and methoxyamine hydrochloride (MeOX), 1% TMCS in *N*-methyl-*N*-trimethylsilyl-trifluoroacetamide (MSTFA), and adonitol, were obtained from Sigma-Aldrich, St. Louis, USA.

### 2.2 Human serum sample

Human serum (Cat. No. H6914, from a male AB clotted whole blood, USA origin, sterile-filtered) was obtained from Sigma-Aldrich, St. Louis, USA.

### 2.3 Serum sample extraction and derivatization for GC-MS and GC-FID analysis

Serum samples (30 μL) were subjected to sequential solvent extraction once each with 1 mL of acetonitrile: isopropanol: water (3:3:2, v/v) ratio and 500 μL of acetonitrile: water (1:1, v/v) ratio mixtures at 4 °C(17). Adonitol (5 μL from 10 mg/ml stock) was added to each aliquot as an internal standard prior to solvent extraction. The pooled extracts (~ 1500 μL) from the two steps were dried under vacuum at 4 °C and parallel extractions performed on empty microcentrifuge tubes only served as extraction blanks to account for background (extraction conditions, derivatization reagents) noise and other sources of contamination (septa, liner, column, vials, handling among others). Blanks were intermittently used to see that no carryovers occurred during randomized run orders and to manually filter out contaminating chemicals from the combined list of features obtained from the blanks. Samples were then sequentially derivatized with methoxyamine hydrochloride (MeOX) and 1% TMCS in *N*-methyl-*N*-trimethylsilyl-trifluoroacetamide (MSTFA) as described elsewhere (7, 18, 19). Steps involved addition of 10 μL of MeOX (20 mg/mL) in pyridine, followed by incubation under shaking at 55 °C for 60 min followed by trimethylsilylation at 60 °C for 60 min after adding 90 μL MSTFA as described (2, 7).

### 2.4 GC-Orbitrap-MS instrument parameters

A robotic arm TriPlus™ RSH autosampler (Thermo Fisher Scientific™, Bremen, Germany) injected 1μL of derivatized sample into a Programmable Temperature Vaporizing (PTV) injector at initial temp of 90 °C to a transfer temp of 290 °C on TRACE™ 1310 gas chromatograph (Thermo Fisher Scientific™, Austin, TX). Helium carrier gas at a flow rate of 1.6 mL/min was used for separation on a Thermo Fisher Scientific™ TG-5MS (60 m length × 0.25 mm i.d. × 0.25 μm film thickness) column. The initial oven temperature was held at 90 °C for 0.5 min, followed by an initial gradient of 10 °C/min ramp rate to 250 °C, where it was held for 5 min, and a gradient of 5 °C/min ramp rate to 295 °C. The final temperature was 295 °C and was held for 35 min. Eluting peaks were transferred through an auxiliary transfer temperature of 250 °C into a Q Exactive™-GC mass spectrometer (Thermo Fisher Scientific™, Bremen, Germany). The mass spectrometer has a resolving power (RP) of 120,000 full width at half maximum (FWHM) at m/z 200 with EI or CI capabilities. From the ion source, an AQT quadrupole is used for precursor ion isolation, which leads into the Orbitrap mass analyzer. Electron ionisation (EI) at 70 eV energy, emission current of 50 μA with an ion source temperature of 230 °C was used in all experiments. A filament delay of 5.3 min was selected to prevent excess reagents from being ionized. High resolution EI spectra were acquired using 60,000 resolution (FWHM at *m/z* 200) with a mass range of *m/z* 50-650.

### 2.5 GC-FID analysis

GC-FID (**Supplementary Figure S1**) analysis was accomplished on the TRACE™ 1310 gas chromatograph (Thermo Fisher Scientific™, Austin, TX). The detector temperature was set at 305 °C where the ignition threshold was 0.5 Pa, airflow of 350 mL/min, hydrogen flow 35 mL/min., and makeup gas 30 mL/min. All other analytic conditions including the column type and column temperature, the injection temperature, splitless injection conditions, carrier gas and the linear velocity were the same as those of GC–MS analysis.

For both analyses, the acquisition sequence started with blank solvent (pyridine) injections, followed by randomized lists of extraction blanks (B), reagent blanks (R), solvent (pyridine-P), and samples (S), where sequences of B, R, and P were injected at scheduled intervals for monitoring shifts in retention indices (RI) as well as serving as system quality control (QC) checks.

### 2.6 GC-Orbitrap-MS data processing

Acquired data was processed using Thermo Fisher Scientific™ TraceFinder™ 4.1 (Thermo Fisher Scientific, Bremen, Germany) software for untargeted analysis. Initial analysis of collected spectra included baseline correction, peak filtering, quantification, assignment of a unique mass and retention indices, signal-to-noise calculation, and compound identification based on the mass spectral pattern as compared to EI spectral libraries. Spectral libraries consulted included: NIST Mass Spectral Reference Library (NIST14/2014; National Institute of Standards and Technology, USA), the Wiley Registry of Mass Spectra – 11^th^ Edition, the MSRI spectral libraries from Golm Metabolome Database (20) available from Max-Planck-Institute for Plant Physiology, Golm, Germany (http://csbdb.mpimp-golm.mpg.de/csbdb/gmd/gmd.html), MassBank (21), MoNA (Mass Bank of North America, (http://mona.fiehnlab.ucdavis.edu/) and a vendor supplied high resolution (HR)-MS mass spectral library for the GC-MS dataset using proprietary TraceFinder™ software (Thermo Fisher Scientific) and MS-DIAL software ver. 3.51 (22) for additional searches, visualization and spectral matching. Further, to filter out noise and less confident compounds, we discarded all compounds with a CV > 30 %. Further, all *siloxane, halogen-derivatives, phthalate, acrylate, and silyloxy, borane, dioxolan,* and *silan, silox,* – derivative compounds were removed from the list manually. For the MS platform, metabolite annotation and assignment followed the metabolomics standards initiative (MSI) guidelines for metabolite identification (23), with Level 2 identification based on spectral database match (match factor >80%) and Level 3 identification where only compound groups were known (specific ions and RT regions of metabolites).

### 2.7 Data sharing

The raw datasets and the metadata obtained from both the platforms are deposited at the Metabolomics Workbench (Study ID: **ST001037**) which are available for download at this link: https://bit.ly/2PlFlW9 (pending publication date).

### 2.8 Statistical analysis

Statistical processing of both GC-FID and GC-MS data sets were performed using statistical software R (Version 3.5.1) (24). Imputed, outlier removed, and scaled peak areas representative of relative metabolite amounts obtained using DeviumWeb (25) are presented. *Univariate and multivariate analysis:* Hierarchical clustering analysis (HCA) was performed on Pearson distances using PermutMatrix (26). The raw metabolite abundance values were Z-score normalized, and the color scale represents +2 (high) to −2 (low) abundance in the heat map. Correlations reported are Pearson correlations which were visualized as heat maps, based on Z-score normalized data ranging from +1 (positive, red), 0 (no correlation, black), and −1 (negative, green) correlation of metabolite abundance across biological and technical replicates. Partial least squared discriminant analyses (PLSDA) were performed using MetaboAnalyst 3.0 (27) and DeviumWeb (25) where the output displayed score plots to visualize the sample groups. The data were scaled with unit variance without any transformation.

### 2.9 Pathway enrichment analysis

Pathway enrichment was performed using MetaboAnalyst 3.0 (www.Metaboanalyst.ca) (27). For ID conversions, the Chemical Translation Service (CTS: http://cts.fiehnlab.ucdavis.edu/conversion/batch) was used in batch mode to convert the common chemical names into their KEGG, HMDB, Metlin, PubChem CID, and ChEBI identifiers.

## 3. Results and Discussion

### 3.1 Comparison of metabolites and peaks from MS and FID detectors

We previously reported on the analysis of non-human primate serum from a baboon using HR-GC-MS alone (7). Here, we expanded our metabolomics analysis to human serum, and compared two orthogonal detection techniques attached to a GC, a MS and a FID detector. Quantitation using a FID is simple, as FID is a mass-sensitive detector that provides a nearly equal molar response to the number of carbon atoms in a hydrocarbon where the detector is fast, and response is linear over a wide dynamic range (~ 10^7^ to 10^8^) (28). A comparison of the FID-chromatogram and total ion chromatogram (TIC) from MS analysis are provided (**Figure 1**). The extracted FID data (filtered) (**Supplementary Table S1**) and the annotated MS-data (**Supplementary Table S2**) are provided. Furthermore, given that there was a solvent delay time used for MS detector, only peaks from 8 to 60 min. (i.e., total 52 min.) were considered for FID peak quantification as well, to only compare peaks/metabolites in affixed retention time windows. We ran five individual serum aliquots (n=5) with three technical replicates each, generating 15 runs and corresponding data files. For MS-based analysis, 2765 metabolites were detected at least once across all the samples (including blanks, reagent blanks, and solvent), which were reduced to 298 compounds that passed all the quality filters described above. However, the S/N criteria used for FID and MS analysis are not comparable as they are different detection methods, and hence, the total number of confident peaks called were very different from both the analysis. For FID-analysis, about 1117 peaks were quantified with retention times across all samples (with < 50% missing values). The median and mean CVs for FID were at 73.43 and 87.90%, whereas those for MS were 43.77 and 51.44%. Thus, with stringent filtering criteria, such as retaining only compounds/ peaks with < 30% RSD, we retained 298 metabolites in the MS analysis and 83 such peaks in the FID analysis. A previous study using fatty acid methyl ester (FAME) analysis showed that 28 FAME standards tested provided similar results for the novel GC-EI-MS-SIM method and GC-EI-MS in the full scan mode, both of which were slightly worse than GC-FID analysis (29).

**Figure 1.**
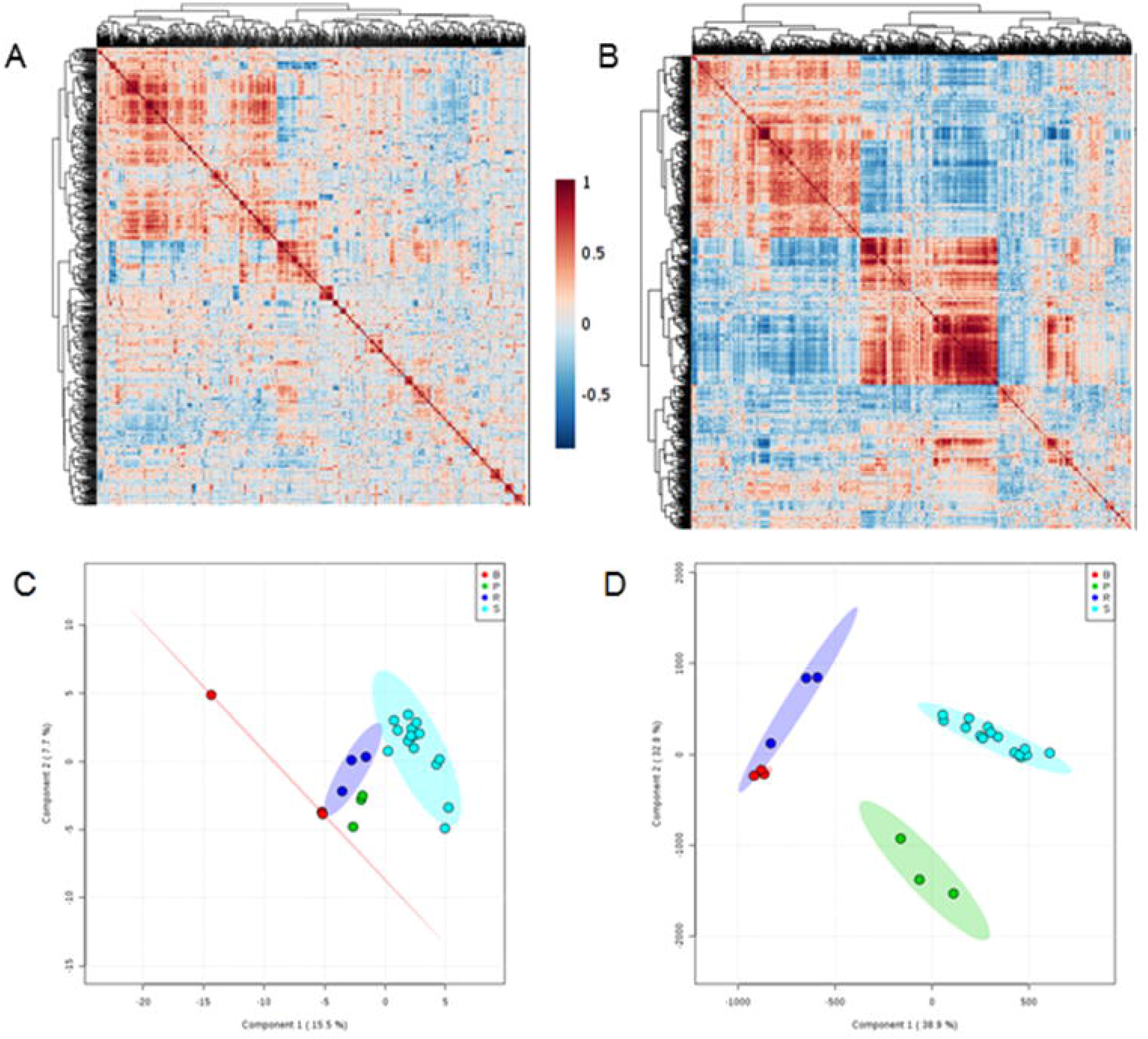
Comparison of chromatograms for human serum sample and representative HR-GC-MS spectra. Chromatograms derived from FID analysis, and a total ion chromatogram obtained from MS analysis, and XIC (m/z 353.17774; unknown) at 5 ppm accuracy are shown. The HRGC-EI-MS spectra of six representative compounds are (A) 2-deoxytetronic acid, (B) methionine, (C) glutamic acid, (D) phenylalanine, (E) lauric acid, and (F) aminomalonic acid.

When we performed hierarchical clustering (HCA) analysis of the top 50 features (either peaks from FID or metabolites abundances from MS) from the two platforms, the results reveal a clearer separation of sample groups (blanks vs samples) for the FID analysis (**Figure 2 A, B**) when compared to those obtained from MS analysis. Similarly, a metabolite-metabolite Pearson correlation analysis for peak and metabolite abundances revealed clearer clusters (two such modules) for the FID data (**Figure 3 A, B**) as opposed to the MS data where the clusters are diffused. When performing a supervised PLS-DA analysis, the FID data explained the clusters better [cumulative score for the first two PCs (PC1, PC2) = 71%] when compared to the MS-data [cumulative score for the first two PCs (PC1, PC2) = 23%] (**Figure 3 C, D**).

**Figure 2.**
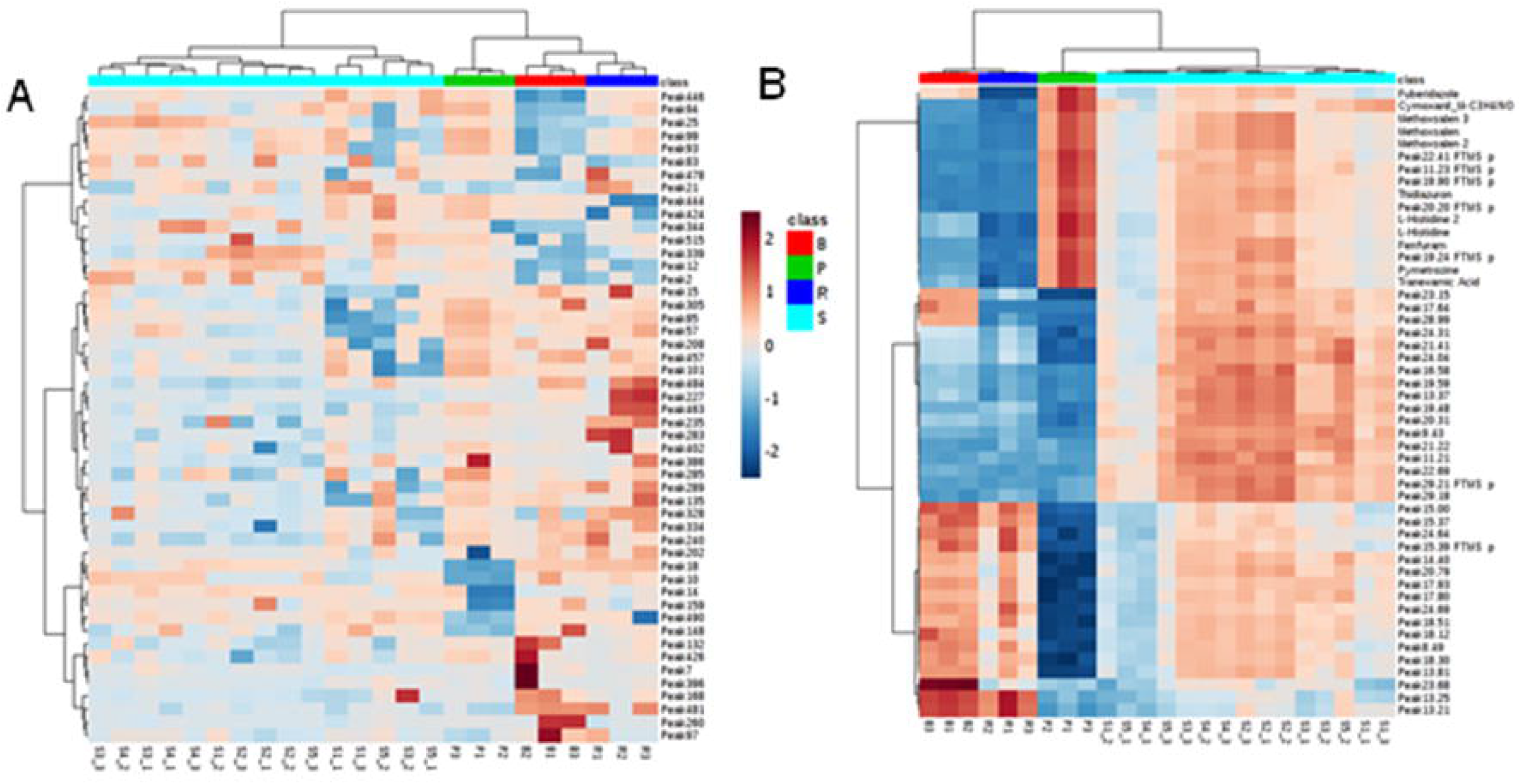
A two-way hierarchical clustering heat map of the serum metabolome (top 50 peaks as obtained from PLS-DA analysis) data for (A) MS detector and (B) FID detector. Each column displays the metabolic pattern of individual samples (extraction blanks, solvents, reagent blanks, and samples). Amount of each peak in individual samples is expressed as relative value obtained by Z-normalization and is represented by the color scheme, in which red and blue indicate high and low concentrations of metabolites, respectively. Rows: samples; Columns: metabolites.

**Figure 3.**
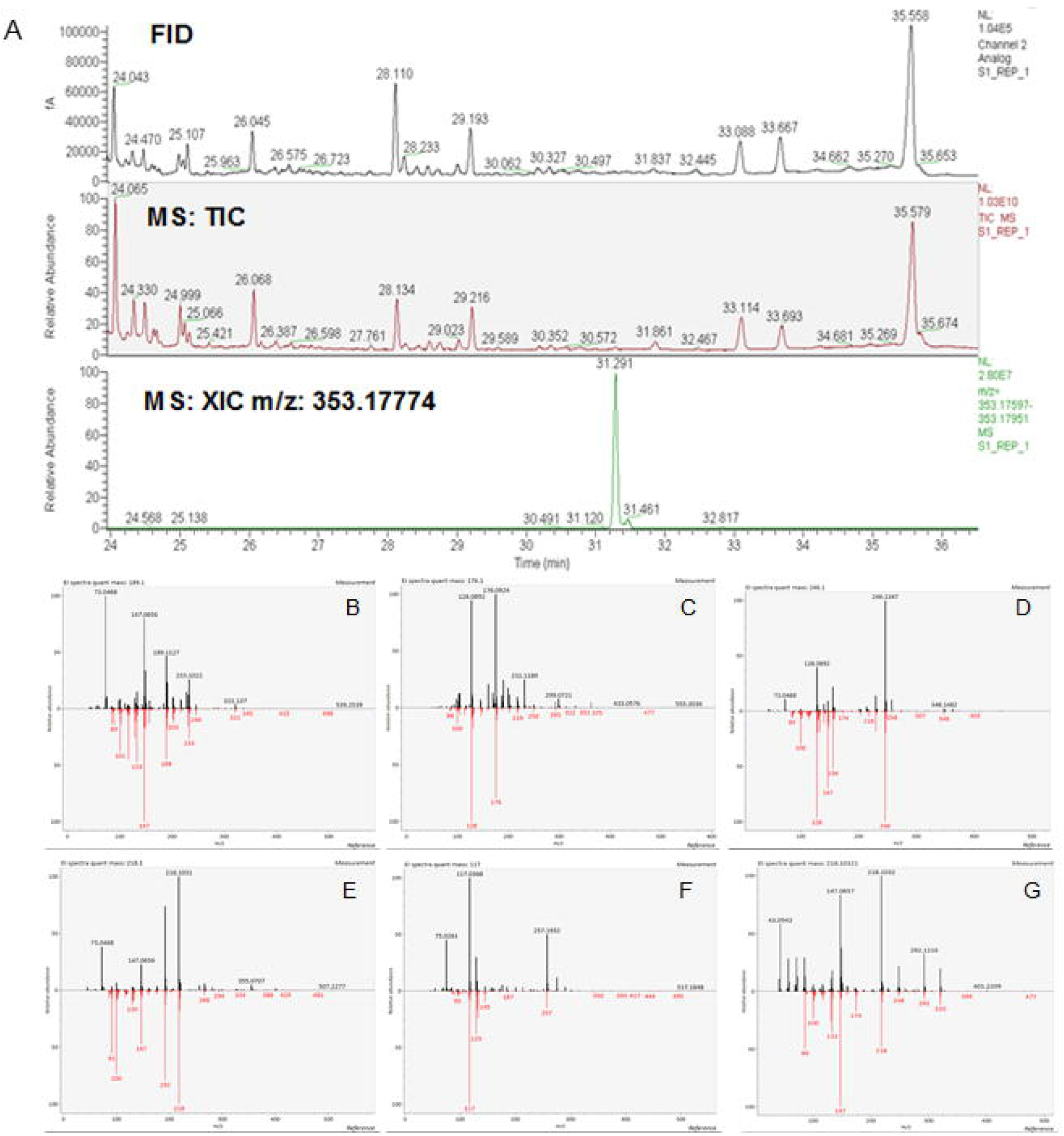
Metabolite-metabolite Pearson correlation map for peak areas for (A) MS analysis and (B) FID-detected peaks. Amount of each peak in individual samples is expressed as relative value obtained by Z-normalization and is represented by the color scheme, in which red and blue indicate high and low values for respectively, for peaks (FID) and metabolites (MS). Supervised PLS-DA analysis for (C) MS analyzed compounds and (D) FID-detected peaks.

We further evaluated the linear correspondence of the quantified compounds based on FID and MS results. We obtained good correlations for randomly handpicked compounds such as glucose, alanine, citric acid, and an unknown, as an example, with correlation scores ranging from 0.99 to 0.89, and fitting linear regression models (**Figure 4 A-D**). Nonetheless, in comparative analysis of volatile compounds in virgin olive oil, it was demonstrated that good selectivity, linearity and higher upper values of the working range are the main advantages of solid-phase microextraction (SPME)-GC-FID versus low bottom values of working ranges, better sensitivity and lower limits of detection and quantification of SPME-GC-MS (30). In another study, no differences associated to particular functional groups were observed between GC-FID and GC-MS, except for the acids, for which working range is much better for GC-FID (30). Also, one-dimensional GC using FID may be sufficient to define biomarker ratios; however, if the samples are too complex, interferences from coeluting compounds will complicate the analysis (31) (Bai et al., 2018).

**Figure 4.**
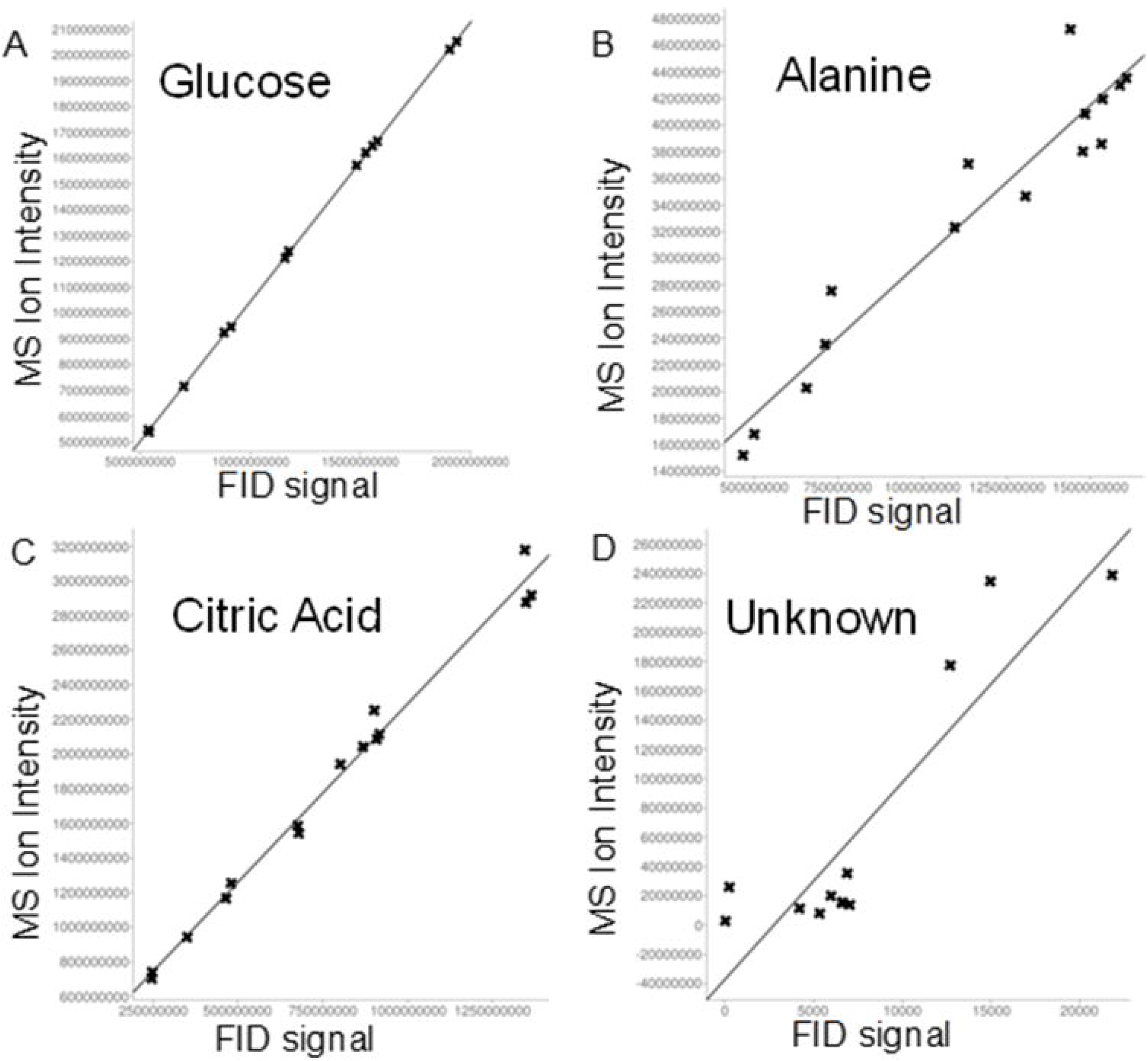
Scatter plots (fitting linear regression models) for FID (x-axis) and MS data (y-axis) for showing linearity in their response factors for all the samples. (A) Glucose [Correlation coefficient (r): 0.9999; Sample size: 13; Intercept (a): −363496920.727; Slope (b): 1.08; Regression line equation: y=1.08x-363496920.727], (B) Alanine (Correlation coefficient (r): 0.958; Sample size 15, Intercept (a): 65761895.608, Slope (b): 0.232, Regression line equation: y=65761895.608+0.232x], (C) Citric acid [Correlation coefficient (r): 0.993; Intercept (a): 229891680.564, Slope (b): 2.059, Regression line equation: y=229891680.564+2.059x) (D) Unknown (Correlation coefficient (r): 0.8954); Sample size 15; Intercept (a): −37477621.483; Slope (b): 13462.916, Regression line equation: y=13462.916x-37477621.483].

### 3.2 Analysis of human serum using FID and HR MS

We analyzed the HRGC-MS data from human serum for its relevance to both clinical and biological analysis. Of the 294 metabolites quantified, we obtained 56 metabolites as trimethylsilylated derivatives and 238 others that were not derivatized (**Supplementary Table S2**). Of these, 133 metabolites were assigned KEGG IDs belonging to various human metabolic pathways. These metabolites included S-adenosyl-L-methionine, adenosine monophosphate, S-adenosyl-L-homocysteine, glucose, alanine, lysine, formic acid, arginine, serine, tryptophan, phenylalanine, urea, 5-phosphorylribose 1-pyrophosphate, biotin, histidine, proline, citric acid, benzoic acid, valine, and threonine. The significantly higher number of metabolites detected in our current efforts, compared to our earlier analysis of a baboon serum sample (7) is attributed to a longer run time of 60 minutes as opposed to the shorter protocol of 23 minutes in the earlier study. Moreover, there are species specific metabolite differences among primate tissues (32) and biofluids. Another 16 metabolites matched KEGG IDs belonging to drugs (i.e., lisinopril, atazanavir, amisulpride, metergoline phenylmethyl ester, alfuzosin decanedioic acid, dibutyl ester, aliskiren, zopiclone, bezafibrate, sulpiride, carbachol, risperidone, ranitidine, indapamide, droperidol). The list also included 41 metabolites that were assigned a LIPIDMAPS ID. These quantified metabolites belonged to 89 various metabolic pathways (and 35 pathways with at least 3 metabolites mapped onto each of them), such as methylhistidine metabolism, thiamine metabolism, glycine and serine metabolism, glucose-alanine cycle, biotin metabolism, carnitine synthesis, transfer of acetyl groups into mitochondria, urea cycle, methionine metabolism, homocysteine degradation, alanine metabolism among others (**Supplementary Table S3**). Recently, GC-FID combined with precolumn derivatization with isobutyl chloroformate was used for confident determination of nucleobases guanine, adenine, cytosine, and thymine from DNA samples (33). Other studies have focused on detection of the food chemicals, i.e., caffeine in coffee grains using GC-FID as well (34). A very recent analysis of a reference material, NIST Standard Reference Material (SRM) 2378 fatty acids in frozen human serum using methods NIST-1 and NIST-2 that use GC-FID and GC-MS platforms, respectively, revealed expanded uncertainties for 12 fatty acids and reference values with expanded uncertainties for an additional 18 fatty acids (35).

Conversely, one cannot map peaks obtained from FID analysis for pathway mapping analysis or enrichment analysis, without access to individual chemical/metabolite standards. However, as mentioned in the previous section, the robust quantification obtained from FID data is advantageous for better quantification when compared to MS-based analysis only. Given that the past FID analysis efforts relied on FAME analysis for metabolite profiling, future analysis can potentially expand on this detection method to use such integrated workflows as the one described in this manuscript. However, robust software tools and analysis workflows that can seamlessly integrate FID and MS-data in real time or offline, are clearly missing.

Nonetheless, both detectors represent a flexible tool for explorative studies and, if supported by appropriate data-processing tools, would appear to be useful in any metabolic profiling study, as was shown using 28 standard compounds including 17 amino acid standards and in CSF samples with simultaneous acquisition with both MS and FID detectors (36). It was also reported that limit of detection (LOD) and limit of quantification (LOQ) are significantly lower for GC-APCI/ToF-MS than for GC-FID. Moreover, the quantitative response of the FID detector is free from ionization bias and those biases introduced by the type of mass analyzer or the instrumental design of a mass spectrometer. Consequently, FID gives a better overall quantitative representation in complex biological samples where traditional MS analysis often is challenged by ion interference effects (36). Further, in a comparison of the non-esterified or free fatty acids quantitative results between the TLC/GC-FID and the GC-MS platforms demonstrated that the GC-MS concentrations of palmitic acid, vaccinic acid, oleic acid, linoleic acid, dihomo-γ-linolenic acid and docosapenta-(4,7,10,13,16)-enoic acid are generally higher than those measured by TLC/GC-FID (11) indicating higher sensitivity and detector-bias as far as MS is concerned.

Our study suffers from several limitations that we clearly recognize, esp. with lower sample size for this proof-of-concept study to demonstrate the applicability of the dual-detector platform for clinical metabolomics studies. Secondly, there are other biofluids such as plasma, saliva and even tissue or cell samples from humans that could be informative for further screening for comparison of those datasets on this new platform. Other complimentary approaches such as high resolution LC-MS/MS or even other detectors such as thermal conductivity detector (TCD) and electron capture detector (ECD) among a host of others would be worth exploring.

## 4. Conclusions

We demonstrated the advantages of a combined GC-FID and HRGC-MS analyses when compared to results obtained from the individual platforms, and how this can boost analytical biochemistry and downstream metabolomics applications. It remains a challenge, like any other untargeted metabolomics platform, to consolidate and align features detected using FID and MS for reliable quantification. We also propose that such instruments which lend the capabilities of detectors that work on different principles would be helpful for correct identification of compounds, especially when standards are available.

## Author contributions

BBM, EB, and MO designed the research; BBM and EB performed the experiments; MO provided essential reagents and materials, BBM and EB analyzed the data, BBM and MO wrote the manuscript, and BBM interpreted the data and has the primary responsibility for the final content and edits. All the authors have accepted responsibility for the entire content of this submitted manuscript and approved submission.

## Human and Animal Rights and Informed Consent

This article does not contain any studies with human or animal subjects performed by any of the authors, but only a de-identified commercially available human serum sample.

## Research funding

None declared.

## Employment or leadership

None declared.

## Honorarium

None declared.

## Conflict of interest

The authors wish to confirm that there are no known conflicts of interest associated with this publication and there has been no significant financial support for this work that could have influenced its outcome.

---

**Supplementary Table S1.** Peak lists obtained for GC-FID analysis.

**Supplementary Table S2**. List of metabolites captured using MS data.

**Supplementary Table S3**. Pathway enrichment analysis for the MS quantified metabolites.

**Supplementary Figure S1**. The GC-FID detector (Thermo Fisher Scientific).

## Notes

https://bit.ly/2PlFlW9

