## Supplementary figures and images for "Comparison of a GC-Orbitrap-MS with Parallel GC-FID Capabilities for Metabolomics of Human Serum"

### Supplementary Figure S1

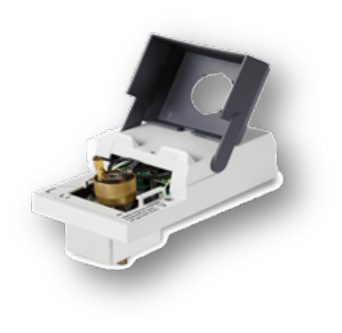

**Supplementary Figure S1.** The GC-FID detector (Thermo Fisher Scientific).
